# Understanding ATP binding to DosS catalytic domain with a short ATP-lid

**DOI:** 10.1101/2023.05.29.542785

**Authors:** Grant Larson, Peter Windsor, Elizabeth Smithwick, Ke Shi, Hideki Aihara, Anoop Rama Damodaran, Ambika Bhagi-Damodaran

**Author notes:** **Corresponding Author** Anoop Rama Damodaran, Ambika Bhagi-Damodaran.

## Abstract

DosS is a heme-sensor histidine kinase that responds to redox-active stimuli in mycobacterial environments by triggering dormancy transformation. Sequence comparison of the catalytic ATP-binding (CA) domain of DosS to other well-studied histidine kinases suggests that it possesses a rather short ATP-lid. This feature has been thought to inhibit DosS kinase activity by blocking ATP binding in the absence of interdomain interactions with the dimerization and histidine phospho-transfer (DHp) domain of full-length DosS. Here, we use a combination of computational modeling, structural biology, and biophysical studies to re-examine ATP-binding modalities in DosS’s CA domain. We show that the closed lid conformation observed in protein crystal structures of DosS CA is caused by the presence of a zinc cation in the ATP binding pocket that coordinates with a glutamate residue on the ATP-lid. Furthermore, circular dichroism (CD) studies and comparisons of DosS CA crystal structure with its AlphaFold model and homologous DesK reveal that a key N-box alpha-helix turn of the ATP pocket manifests as a random coil in the zinc-coordinated protein crystal structure. We note that this closed lid conformation and the random-coil transformation of an N-box alpha-helix turn are artifacts arising from the millimolar zinc concentration used in DosS CA crystallization conditions. In contrast, in the absence of zinc, we find that the short ATP-lid of DosS CA has significant conformational flexibility and can bind ATP (*K*_d_ = 53 ± 13 μM). We conclude that DosS CA is almost always bound to ATP under physiological conditions (1-5 mM ATP, sub-nanomolar free zinc) in the bacterial environment. Our findings elucidate the conformational adaptability of the short ATP-lid, its relevance to ATP binding in DosS CA and provide insights that extends to 2988 homologous bacterial proteins containing such ATP-lids.

## INTRODUCTION

Bacterial species often utilize a two-component signal transduction (TCS) pathway to sense and respond to changes in their microenvironment. This pathway consists of a histidine kinase (HK) and a response regulator (RR) that interact to transfer the signal from the extracellular stimuli to bacterial DNA (**Fig. S1**).^1^ HKs are homodimeric transmembrane enzymes, which upon binding ATP, catalyze auto-phosphorylation of a conserved histidine found in its dimerization and histidine phosphotransfer (DHp) domain in response to the stimuli (**Fig. 1a**).^2^ The phosphoryl group on the HK’s catalytically-activated histidine is then transferred to an aspartate residue of the RR, which binds to bacterial DNA to regulate gene transcription in response to the extracellular stimuli (**Fig. S1**).^3^ As a result, HKs are essential for bacteria to adapt to various environmental changes, making them attractive drug targets due to their orthogonality to eukaryotic sensory domains and their importance to pathogenicity and bacterial survival.^4,5^ One such extensively investigated HK drug target is DosS from *Mycobacterium tuberculosis* (*Mtb*) which triggers dormancy in the bacteria under hypoxic/NO/CO-rich extracellular redox environments, making it resistant to tuberculosis drug regimes.^6–11^ Specifically, upon sensing hypoxia or nanomolar amounts of NO/CO, DosS catalyzes auto-phosphorylation of its H395 residue and then transfers the phosphoryl group to the D54 residue of the RR, DosR. The phosphorylated DosR, in turn, activates the transcription of more than 50 dormancy-related genes in *Mtb*, including several genes implicated in the proteostasis network.^12,13^ Despite consensus on downstream phosphotransfer events, the molecular basis for controlling interdomain redox signal transduction within the DosS heme sensor system remains unclear.^14,15^ Previous studies have suggested that the short ATP-lid in the catalytic ATP-binding domain (CA) of DosS inhibits kinase activity by blocking ATP binding in the absence of interdomain interactions with the DHp domain of full-length DosS.^16^ Given the significance of DosS in *Mtb*’s persistence and drug resistance, it is critical to closely examine and understand DosS’s ATP-binding modalities in comparison to other well-studied HKs.

**Figure 1.**
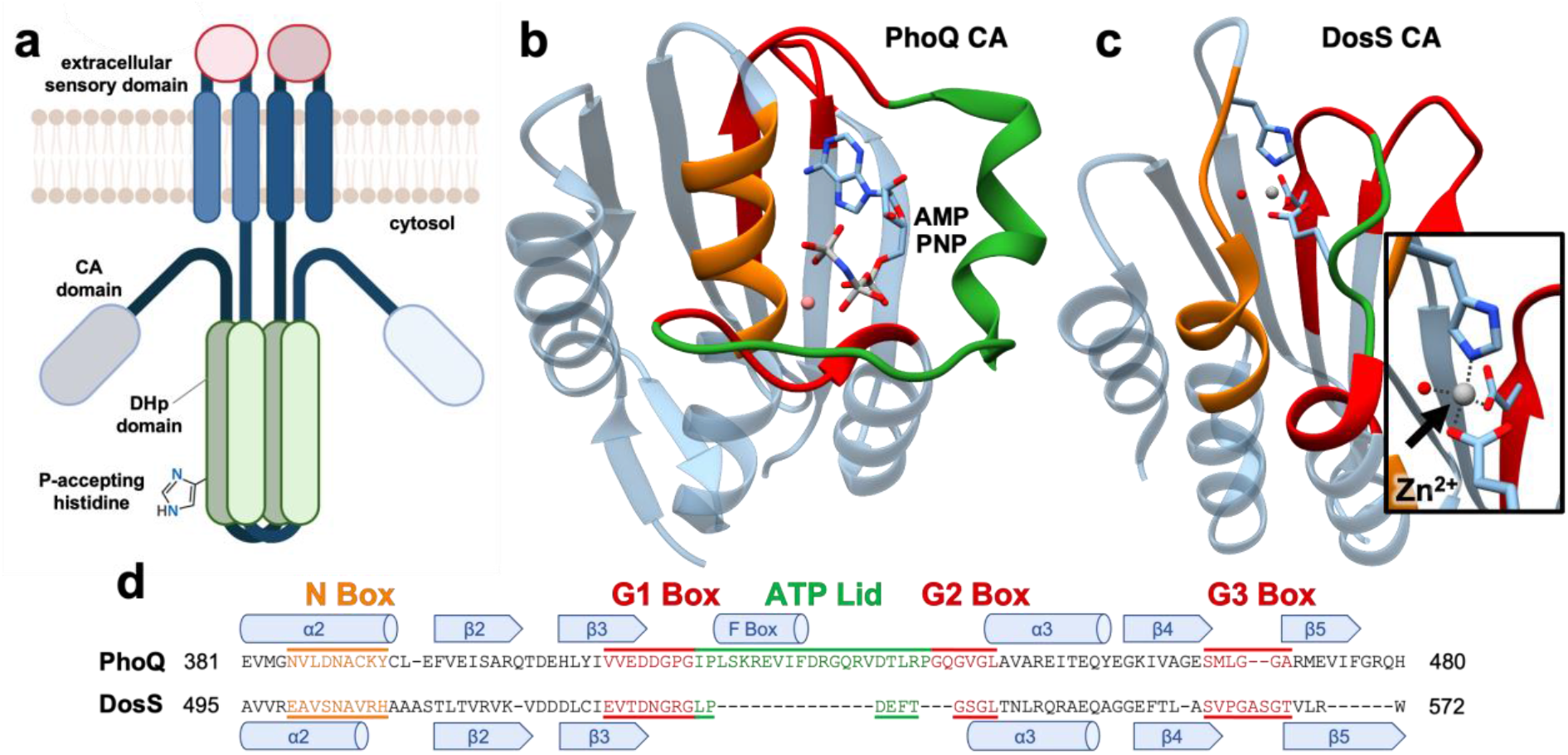
**a)** Representation of a dimeric histidine kinase (HK) complex. A sensory domain (in pink) is located in the extracellular space while the dimerization and histidine phosphotransfer (DHp) domain (in green) and the catalytic ATP-binding domain (CA in grey) extend into the intracellular cytosolic space. Crystal structures of (b) the PhoQ CA bound to an ATP analog (AMPPNP; PDB ID: 1ID0) and (**c**) DosS CA bound to a zinc ion (PDB ID: 3ZXO). Zinc coordinating residues are highlighted in the inset of panel c. In both structures, the N-box is colored orange, G-boxes are colored red, and the ATP-lid is colored green, **d**) Alignment of DosS and PhoQ CA protein sequences showing various boxes and secondary structures.

The ATP-binding pocket of a typical HK CA is an α-helix/β-sheet sandwich (the Bergerat fold) featuring several conserved amino acid sequences referred to as “boxes”.^17^ How these boxes contribute to ATP binding is well understood for HKs like PhoQ and DesK.^18,19^ For instance, the PhoQ CA (**Fig. 1b**) contains one N-box (colored orange), three G-boxes (colored red), and one F-box containing ATP-lid (colored green). The N-box has conserved asparagine, histidine, and lysine/arginine residues that bind to the triphosphate chain of ATP via ionic and hydrogen bonding (H-bonding) interactions. All three G-boxes possess multiple glycine residues that afford flexibility to the protein’s macromolecular structure for optimal ATP binding.^20^ Moreover, the adenine moiety of ATP is H-bonded to a conserved aspartate in the G1-box. Additionally, the CA domain comprises an ATP-lid that is characterized by considerable conformational flexibility in the absence of any bound nucleotide, and often includes a conserved phenylalanine containing F-box. Upon nucleotide binding, the lid interacts with the DHp domain to correctly position the γ-phosphate of ATP for an autophosphorylation reaction.^17^ However, the ATP-lid displays length and secondary-structure variations across different HK types. For instance, the ATP-lid of PhoQ CA has 21 residues and is significantly longer than that of DosS which has only 6 residues with the F-box absent (highlighted green in **Fig. 1c-d, Fig. S2**). This short ATP-lid motif has been hypothesized to modulate DosS kinase activity by preventing ATP binding in the absence of interdomain interactions with the DHp domain of full-length DosS.^16^ In this work, we use a combination of computational modeling, structural biology, and biophysical investigations to further explore the structural and molecular aspects of ATP binding to the DosS CA domain. We show that despite possessing a short ATP lid, DosS CA exhibits an affinity to ATP (*K*_d_ = 53 ± 13 μM) that is comparable to other HK’s and is usually bound to ATP under physiological conditions of 1-5 mM ATP in the bacteria.^21^ Overall, our studies demonstrate key structural aspects and molecular interactions that enable ATP binding to DosS and are applicable to 2988 other homologous bacterial kinases that feature a short ATP-lid.

## MATERIALS AND METHODS

All reagents were purchased from commercial sources and used without further purification.

### Plasmid design and mutagenesis

pET23a(+) vector encoding WT DosS CA (amino acids 454 – 578 of full-length DosS) sequence with an N-terminal 6xHis tag and a TEV protease site was purchased from GenScript. DosS CA mutants were prepared from the WT plasmid via site-directed mutagenesis with end-to-end primers similar to methods described in previous papers.^22^ The forward and reverse primers (5’ to 3’) for the H507F mutation were: ACACTTACAGTAAGAGTGAAGGTAGACGATGATTTG and GGATGCCTTCGCGAATC-TAACAGCGTT, respectively. The forward and reverse primers (5’ to 3’) for the D529N mutation were: TCGAAGTCACTAATAACGGGCGGGG and TGCACAAATCATCGTC-TACCTTCACTCTTACTG, respectively. The forward and reverse primers (5’ to 3’) for the E537A mutation were: GTTACCTGATGCGTTCACTGGCTCCG and CCCCGCCCGTTATCAGTGACTTCGAT, respectively. All plasmid sequences were confirmed via sequencing by ACTG.

### Expression and purification of WT DosS CA and its variants

BL21(DE3) competent cells (ThermoFisher) were transformed with the pET23a(+)-CA plasmid and plated on LB agar plates containing 100 μg/mL ampicillin. Primary cultures were inoculated with a single colony from an agar plate and were grown overnight at 37 °C while shaking at 220 rpm. The secondary cultures were inoculated with 25 mL of primary culture and were grown in 1.5 L 2XYT media containing 100 μg/mL ampicillin at 37 °C while shaking at 220 rpm. The cell density was monitored until the OD_600_ reached 0.6 – 0.8. Protein expression was induced with 0.5 mM isopropyl 1-thio-□-D-galactopyranoside (IPTG) at 18 °C while shaking at 180 rpm for 18 hours. Cells were harvested via centrifugation at 8000 rpm for 15 min at 4 °C. The cell pellet obtained was re-suspended in 50 mM Tris (pH 7.5), 250 mM sodium chloride, and 1% Triton X-100 and lysed via sonification (Fisher Scientific) while on ice. The crude lysate was centrifuged for 1 hour at 20,000 rpm and 4 °C (Beckman Coulter). The supernatant was filtered using a 0.44 μm syringe filter (Sartorius) and applied to a 5 mL HisTrap FF column (GE Healthcare) at a rate of 2 mL/min. The column was washed with 75 mL 50 mM Tris (pH 7.5), 250 mM sodium chloride, and 20 mM imidazole at a rate of 2 mL/min. Recombinant protein was eluted with 50 mM Tris (pH 7.5), 250 mM sodium chloride, and 250 mM imidazole. The eluted protein was concentrated and buffer exchanged into 50 mM HEPES (pH 7.5), 15 mM MgCl_2_, 94.5 mM NaCl, and 5% glycerol. Finally, the protein solution was concentrated to ∼6 mL using a centricon (MW cutoff = 10 kDa) and was purified via size exclusion chromatography with a HiLoad 26/600 Superdex 75 pg column (GE Healthcare). The purified protein solution was diluted to 135 μM (∼2.0 mg/mL) in buffer containing 0.5 mM EDTA and 1 mM DTT, and 4.5 μM TEV Protease was added to it. The reaction mixture was gently rotated overnight and cleaved protein product was purified from the uncleaved protein using a HisTrap FF column. EDTA and DTT were removed via a PD-10 column (GE Healthcare) equilibrated with 50 mM HEPES (pH 7.5), 15 mM MgCl_2_, 94.5 mM NaCl, and 5% glycerol. The protein solution was concentrated to 2.25 mM, as estimated on the basis of absorbance at 280 nm using the extinction coefficient 5579 M^-1^ cm^-1^ determined via BCA assay. Variants of DosS CA were expressed and purified using a similar protocol with minor variations. The purity of DosS CA protein samples were verified using SDS PAGE gel (**Fig. S3**).

### X-ray crystallography and protein structure determination

Initial crystallographic screening used sitting-drop vapor diffusion method, conducted at the Nanoliter Crystallization Facility at the University of Minnesota (Minneapolis, MN). For each drop, 100 nL of 581 mM ABD (7.7 mg/mL) was mixed with 100 nL of commercial screening kits, namely Index (Hampton Research), PEG/ION HT (Hampton Research), and Shotgun1 (Molecular Dimensions). A total of 288 different conditions were tested, 864 total drops with three samples per condition. Diffraction-quality crystals were generated in 24-well hanging drop crystallization plates by mixing 1-2 μL of 581 μM CA (7.7 mg/mL) with 1-2 μL of 10 mM ZnCl_2_, 20% PEG 6000 (w/v), and 100 mM MES (pH 6.0) or 100 mM HEPES (pH 7.2). The CA crystals were cryo-protected by soaking in the reservoir solution supplemented with ethylene glycol, by gradually increasing ethylene glycol concentration to 20%, and subsequently flash frozen in liquid nitrogen. X-ray diffraction data were collected at the NE-CAT beamline 24-ID-C of the Advanced Photon Source (Lemont, IL) and processed using XDS.^23^ The structures were determined by molecular replacement with PHASER using DosS (PDB ID: 3ZXO) as the search model.^16,24–27^ Iterative model building and refinement were conducted using COOT and PHENIX.^28,29^ A summary of crystallographic data statistics is shown in **Table S1**. Figures were generated using Chimera.^30^ The coordinates and reflection files were deposited with rcsb.org with the accession code 8SBM.

### Molecular Dynamics (MD) studies

For the simulation of ATP-lid dynamics the starting structure of WT DosS CA was taken from the crystal structure (PDB: 8SBM). Using the tleap module in AmberTools22, explicit hydrogen atoms were added, Na^+^ ions were added to a concentration of 0.15 M, the system was neutralized with Cl^-^ counter ions, and the DosS CA protein was parameterized using the ff14SB forcefield.^31,32^ For simulations including Zn^2+^, a forcefield modification was included as well as a restraint that maintained the following distances: ∼1.94 Å between Zn^2+^ and H507, ∼1.82 Å between Zn^2+^ and D529, ∼2.00 Å between Zn^2+^ and E537, and ∼1.95 Å between Zn^2+^ and a water molecule.^33^ The protein was solvated in a 10.0 Å cuboid unit cell with TIP3P water molecules.^34,35^ The protein system was minimized, gently heated to 300, and the density was equilibrated for 2 ns. Three independent 300 ns simulations were performed. CPPTRAJ was used for to perform all trajectory analyses. To simulate the protein-nucleotide interactions of the ATP-docked DosS CA structure, atomic coordinates were obtained from the results of a docking calculation. The system was prepared using AmberTools22 as described above.^31^ In addition to the ff14SB forcefield, the DNA.OL15 forcefield was applied to parameterize ATP, and a forcefield modification was applied to the polyphosphate group.^36,37^ Restraints were placed between atom N6 of ATP and a carboxylate oxygen atom of D529 to maintain a distance ∼3.0 Å, Mg^2+^ and an oxygen atom on the γ-phosphate of ATP, and Mg^2+^ and an oxygen atom on the β-phosphate of ATP to prevent dissociation. The distance of the Mg^2+^ restraints was centered at 2.0 Å. The system preparation, minimization, equilibration, and heating were performed as described above. Three independent 300 ns simulations were performed. CPPTRAJ was used for to perform all trajectory analyses.

In the DosS CA Alphafold model simulations, the atomic coordinates of DosS CA were obtained from the AlphaFold predicted structure of DosS.^38^ The DosS CA structure was aligned to the ATP-bound crystal structure of the homologous DesK CA domain (PDB: 3EHG) using the MatchMaker tool in Chimera.^39^ The rigid body superposition of DosS and DesK allowed for the atomic coordinates of ATP from DesK to be superimposed onto the simulated DosS CA domain structures.^40^ The system was prepared using AmberTools22 as described above.^31^ In addition to the ff14SB forcefield, the DNA.OL15 forcefield was used to parameterize ATP, and a forcefield modification was applied to the polyphosphate group.^36,37^ System preparation, minimization, equilibration, and heating were performed as described above. No restraint was needed to maintain ATP interactions with DosS CA. Three independent 300 ns simulations were performed. CPPTRAJ was used for to perform all trajectory analyses.

### Docking Studies

Atomic coordinates of the DosS CA were obtained from frame 493 of the molecular dynamics simulation above in which the ATP-lid motif formed an open conformation, which was determined by finding the maximum distance between the C_α_’s of His507 and Glu537 (**Fig. 2c**). Molecular docking simulations were performed using Glide (Schrödinger). The protein structure was minimized using default parameters. We constructed ATP-Mg^2+^ with Mg^2+^ bound to oxygen atoms on the γ- and β-phosphate groups and minimized the complex using default parameters. The receptor grid was generated and centered around Asp529, a conserved residue that forms analogous interactions between ATP and homologous CAs. Residues surrounding the presumed binding pocket of ATP were allowed to rotate freely during the docking simulation. ATP-Mg^2+^ was docked to the protein structure using Glide with flexible ligand sampling, XP scoring function, and post-docking minimization. All other parameters were kept as default.

**Figure 2.**
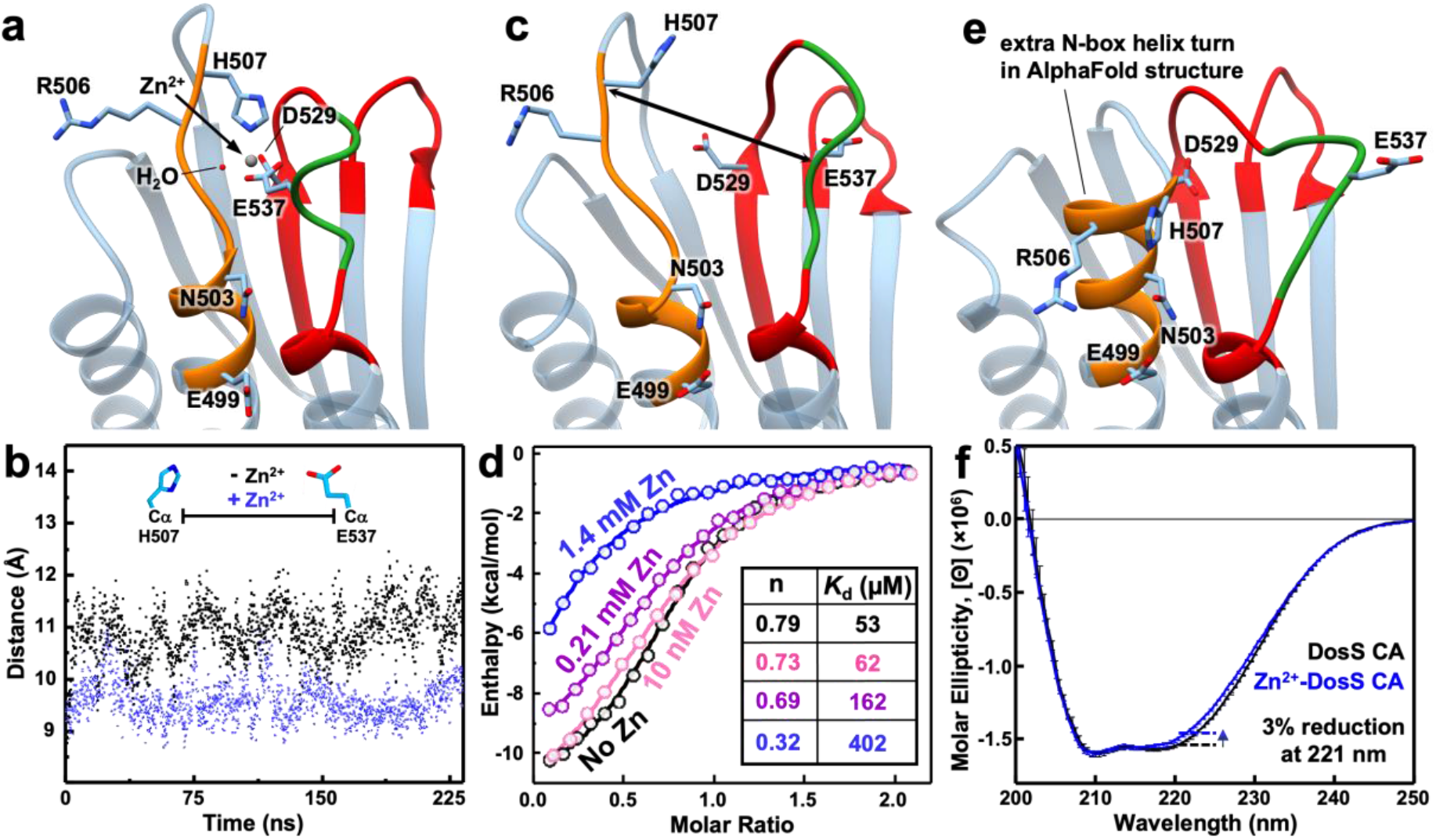
**a)** Crystal structure of DosS CA showing the primary coordination sphere of zinc (PDB ID: 8SBM). **b**) Distance variation between alpha carbons of H507 and E537 residues of DosS CA with zinc (blue) and without zinc (black) during MD simulations, **c**) Structure of the MD frame with the greatest distance between H507 and E537. **d**) ITC data with DosS CA in the presence of 0 pM (black), 10 nM (pink), 0.21 mM (purple), and 1.4 mM (blue) zinc. . A table with Kd and n value for these variants is summarized in the inset, **e**) AlphaFold structure of the DosS CA domain shows an extra alpha-helix in the N-box. **f**) CD data of DosS CA in the absence (black) and presence (blue) of 10 mM zinc. Inset shows zoomed-in CD data in the 198-210 nm wavelength range.

### Isothermal Calorimetry (ITC) Measurements

ITC experiments were performed using a low volume Nano ITC (TA Instruments) similar to previous studies.^41,42^ A 0.5 mL 600 μM protein solution was prepared and placed in 6 kDa molecular weight cutoff dialysis membrane (Spectra/Por). The protein solution was dialyzed overnight against 50 mM HEPES (pH 7.5), 15 mM MgCl_2_, and 94.5 mM NaCl while slowly stirring at 4 °C. This protein solution was filtered via 0.22 μm syringe filter and then used to prepare a 400 μL 400 μM protein 50 mM HEPES (pH 7.5), 15 mM MgCl_2_, 94.5 mM NaCl, and 0.8 mM TCEP titrand solution. A 400 μL 2.4 mM AMP-PNP solution containing 0.8 mM TCEP was prepared using filtered dialysate. A control titrand solution containing no protein and 0.8 mM TCEP was prepared with filtered dialysate. The sample cell was filled with 300 μL of protein or control solution and was cooled to 10 °C. The syringe was loaded with 50 μL of nucleotide solution. The nucleotide solution was titrated into the sample cell with one 0.5 μL and twenty-four 2 μL injections while stirring at 200 rpm with 200 seconds in between injections to ensure thorough mixing of the cell. Data analysis was performed using the NanoAnalyze software (TA Instruments). The enthalpy of dilution of nucleotide was subtracted from the raw data. Corrected heat data were integrated peak by peak and the area was normalized per mole of injectant to get a plot of observed enthalpy change per mole of nucleotide against the molar ratio of nucleotide to protein.

### Circular Dichroism of DosS CA

Solutions of 40 μM DosS CA were prepared with and without 10 mM Zn(OAc)_2_ in buffer containing 10 mM BisTris pH 7 and 50 mM NaF. A 1 mm quartz cuvette was used for sample measurement in a Jasco 815 Circular Dichroism spectrometer. Samples were measured between the wavelengths of 190 to 260 nm at 0.5 nm intervals with a bandwidth = 2 nm, data integration time = 2 seconds, and a scan rate = 50 nm/min. A baseline was obtained using buffer containing 10 mM BisTris pH 7 and 50 mM NaF with and without 10 mM Zn(OAc)_2_. Ellipticity (Θ, mdeg) was converted to Molar Ellipticity ([Θ], (deg * cm^2^) / dmol) using the following equation [Θ] = (Θ * MW) / (PL * C * 10), where MW is the Average Molecular Weight, PL is the pathlength (in cm), and C is the protein concentration (in mg/mL). The calibration of the instrument was validated with a 0.06% CSA solution.

### Bioinformatics studies

A sequence-based homology search (NCBI BLAST) of residues 454 – 578, the DosS CA domain, was performed using the non-redundant protein sequences database excluding Mycobacterium (taxid: 1763) organisms (BLAST E-value of e^-5^).^43^ The sequences of related CA domains were aligned using the NCBI Constraint-based Multiple Alignment Tool (COBALT).^44^ Next, CA domains were sorted based on the number of amino acid residues in the ATP-lid/F-box region, which was defined as the amino acids between the G1- and G2-box. Finally, a sequence logo plot was generated using WebLogo to analyze the conservation of amino acid residues in the homology boxes of CA domain sequences.^45^

## RESULTS AND DISCUSSION

To obtain a molecular understanding of the ATP-binding site in DosS, we performed a shotgun screen and identified crystallization conditions for WT DosS CA. Consistent with previous structural studies on a DosS CA variant,^16^ successful crystallization of WT DosS CA required 10 mM zinc, and the resulting protein crystals diffracted at 1.5 Å resolution. The WT DosS CA structure shows a zinc ion in the ATP binding pocket coordinated to H507, D529, and E537 residues as well as a water molecule H-bonded to the carbonyl oxygen atom of V505 in a distorted tetrahedral geometry (**Fig. 2a**). Among these, D529 is a highly conserved aspartate residue in the G1-box and is considered as the primary site of interaction between HKs and ATP’s N6 amino group. At the same time, zinc coordination to E537 residue in the ATP-lid may hinder conformational restructuring required for ATP binding. To investigate such effects, we performed MD simulations of the DosS CA in the presence and absence of zinc. We tracked distance variation between the C_α_’s of H507 and E537 residues throughout the course of the MD simulations. Residues H507 and E537 are present on loops 1 and 2 in the DosS CA structure (**Fig. S4a**), and the distance between their C_α_’s provides a measure of the openness of the ATP-lid. Temporal variations of this distance also provide a measure of ATP-lid flexibility. The C_α_ distance stays nearly constant at 9-10 Å for the zinc bound CA structure suggesting low conformational flexibility of the protein and a relatively closed pocket in the presence of zinc (**Fig. 2b** blue trace). However, when zinc is removed, we observe that the C_α_ distance rapidly increases up to 12 Å and varies from 12.5 Å to 9.5 Å over the 300 ns simulation time suggesting a significantly open ATP pocket and enhanced lid flexibility (**Fig. 2b** black trace and **Fig. 2c**).

In order to experimentally probe the influence of zinc on ATP-lid flexibility and ATP binding to DosS CA, we conducted detailed isothermal calorimetry (ITC)-based binding affinity studies of a non-hydrolysable ATP analog, adenylyl-imidodiphoshate (AMP-PNP) ^46^ to DosS CA. We began by titrating AMP-PNP to WT DosS CA in the absence of any zinc and observed an exothermic heat profile indicative of an enthalpically driven binding interaction between the nucleotide and DosS CA (**Fig. S5**). Using a single site binding equation to fit the heat evolved per mole of added ligand *vs* ligand-to-protein molar ratio, we extracted various parameters including binding affinity (*K*_d_ = 53 ± 13 μM), goodness of fit (*c* value = 6.2), maximal heat released (*ΔH*_*max*_ = -12.4 kcal/mol), and binding stoichiometry (*n* = 0.79) for the interaction between DosS CA and AMP-PNP. The *K*_d_ value of 53 ± 13 μM for AMP-PNP binding to DosS CA is comparable to other HKs such as HK853 and Spo0B (**Table S2**).^47,48^ This suggests that DosS CA would be completely bound to ATP under physiologically relevant ATP concentrations (1-5 mM) in bacteria and confutes claims from previous work that DosS CA by itself is unable to bind ATP.^16,21^ To probe the influence of zinc on nucleotide binding to DosS CA, we also conducted ITC titrations of AMP-PNP to DosS CA in the presence of varied zinc concentrations. Cells typically have all of their zinc tightly bound to proteins and other cofactors with only femtomolar concentrations of free zinc.^49,50^ Our results show that even at 10 nM zinc concentrations, the binding of AMP-PNP to DosS CA remains largely unaltered (*K*_d_ = 62 μM, n = 0.73) indicating negligible effect (**Fig. S6**). We only observed competitive inhibition of AMP-PNP binding to DosS CA at exceedingly high zinc concentrations of 210 μM (**Fig. 2d** purple curve, *K*_d_ = 162 μM, n = 0.69) and 1.4 mM (**Fig. 2d** blue curve, *K*_d_ = 402 μM, n = 0.32) (**Fig. S7**). In addition to the diminished binding affinity, a significant decrease in binding stoichiometry is indicative of progressive enzyme deactivation at millimolar zinc concentrations, which are relevant to WT DosS CA crystallization conditions. A closer examination of the DosS CA structure crystallized under millimolar zinc concentrations with its AlphaFold model (**Fig. 2e**) and its DesK homolog (**Fig. S8**) reveals that a key N-box alpha-helix turn (residues N503-H507) of the ATP pocket manifests as a random coil in the zinc-coordinated protein crystal structure. Circular Dichroism (CD) can be used to detect such changes in protein secondary structure. In particular, helix-to-coil transitions have been extensively studied using CD and are characterized by a reduction in the alpha-helix-related dip at 221 nm.^51^ We conducted comprehensive CD studies of DosS CA under zinc-free and 10 mM zinc conditions (**Fig. 2f** black and blue curves, respectively). As anticipated, we observe signatures consistent with a helix-to-coil transition in the presence of zinc highlighted by a ∼3% reduction in the dip at 221 nm. Although these are small yet detectable changes in CD spectrum, the magnitude correlates with the random coil transition of a single turn of an alpha helix in DosS CA, which constitutes ∼7% of helical content in the protein (**Fig. S4**). These findings suggest that the tetrahedral coordination of zinc to residues in the ATP pocket, and more specifically to H507, drives a random coil transition of a key N-box helix turn, presenting itself as an artifact in the protein crystal structure that is physiologically irrelevant. In conclusion, these studies demonstrate that the ATP-lid of DosS CA has significant flexibility, ATP can effectively bind to DosS CA, and that the AlphaFold model likely presents a physiologically relevant representation of DosS CA as compared to the crystallized zinc-bound structure.

Next, we performed MD simulations on the AlphaFold model of DosS CA using ATP coordinates taken from its superposition to the homologous ATP-bound DesK crystal structure.^2^ The simulations showed that ATP remained in DosS CA’s ATP pocket for >100 ns of simulation time (**Fig. 3a**) and its placement in the pocket resembled that of ATP-bound crystal structures of PhoQ and DesK (**Fig. S9**). The simulated structure captured several characteristic interactions between ATP and residues in the G- and N-boxes of DosS CA (**Fig. 3a**). For instance, D529 in the G1-box and T568 in the G3-box interacted with ATP’s adenine ring. D529 formed an H-bond to N6 of adenine and, along with T568, positioned a water molecule to H-bond with N1 of adenine. Furthermore, S541 in the G2-box formed an H-bond with the γ-phosphate. Moving to the N-box, H507 formed H-bonds to the oxo groups of the α- and β-phosphates while R506 formed H-bonds to the oxo groups of the β- and γ-phosphates. Additionally, the carboxylate of E499 and the carbonyl of N503 also coordinated to the Mg^2+^ cation associated with ATP. We found no interactions between ATP and residues on the ATP-lid residues. We compared these observations with simulations of ATP docked to the zinc-removed DosS CA crystal structure and found that the random coil transition of an N-box alpha helix turn results in an unusual (out-of-pocket) placement of ATP that only interacts with D529 and R506 residues (**Fig. S10)**. To further assess the physiologically relevant mode of ATP binding to DosS CA, we conducted detailed ITC-based binding affinity studies of AMP-PNP to DosS CA variants with selected residues mutated (**Fig. S11**). We began with the E537A variant, in which the zinc-coordinating glutamate on the ATP-lid that is responsible for the closed-lid crystal structure was mutated to an alanine. ITC studies revealed only a slightly weakened binding affinity of 96 μM (**Fig. 3c**, green curve) compared to WT DosS CA (**Fig. 3c**, black curve). The lack of any affinity enhancement despite mutation of the ATP-lid residue responsible for the closed state in the presence of zinc further confirms that the short ATP-lid does not hinder ATP binding. Moving on to the D529N variant (**Fig. 3c**, red curve), which corresponds to a mutation of the adenine-binding aspartate in the G1-box to an asparagine, we observed complete loss of AMP-PNP binding capacity. This is expected, as D529 is a highly conserved aspartate residue that binds to the adenine moiety of ATP across different HK types as observed in both our MD simulations which utilized the AlphaFold structure (**Fig. 3b**) and the zinc-removed crystal structure (**Fig. S10**). Focusing on the H507F variant which corresponds to a mutation of the N-box histidine to phenylalanine, we observed a *16*-fold weakening of AMP-PNP binding affinity (**Fig. 3c**, orange curve) confirming the N-box H507 residue to be important for ATP binding. We note that H507 formed an H-bond with the oxo groups of the α- and β-phosphates in MD simulations that employed the AlphaFold model of DosS CA (**Fig. 3b**) but was non-interacting in MD simulations that employed the DosS CA crystal structure (**Fig. S10**). We believe that the non-interaction is due to the random coil manifestation of the N-box helix turn containing residues N503-H507, which orients the residues in such a way that only R506 can participate in ATP binding. Consequently, these ITC studies further validate the MD-based ATP-binding pose obtained using the AlphaFold model and highlight artifacts in the zinc-containing crystal structure. Overall, our computational modeling and biophysical studies demonstrate that the ATP pocket in DosS CA can bind ATP despite its short ATP-lid.

**Figure 3.**
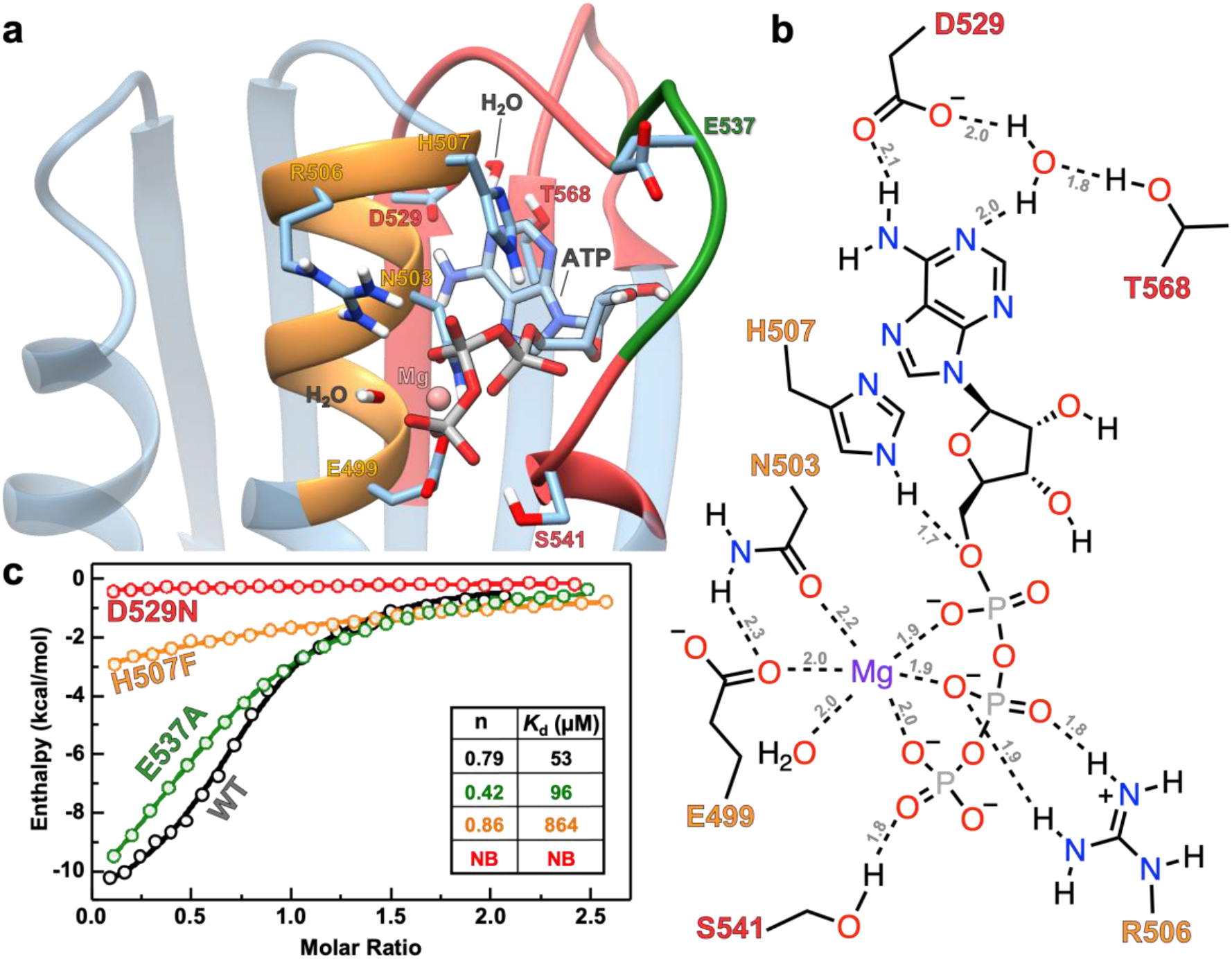
**a)** MD simulated structure of ATP bound to AlphaFold model of DosS CA. **b)** Schematic showing molecular interactions between DosS CA residues and ATP from MD simulations. H-bonds are indicated with dashed lines and the distances (in Angstroms) are labeled in grey. Bond distances and angles are not to scale. The color of residue labels in (a) and (b) match that of the boxes to which they belong. Orange is for N-box, red is for G-box, and green is for the ATP-lid. **c)** ITC titrations showing ATP analogue (AMP-PNP) binding affinity for WT (black), E537A (green), D529N (red), and H507F (orange) DosS CA variants. A table with *K*_d_ and n value for these variants is summarized in the inset. No binding is abbreviated as NB.

Having demonstrated that DosS CA is capable of binding ATP with an affinity comparable to prototypical HKs with F-boxes and long ATP-lids, we set out to identify if other DosS-like proteins that lack an F-box and possess shorter ATP-lids are widespread. To this end, we searched the NCBI database for non-redundant proteins and found 5000 proteins homologous to DosS CA. We chose top 3000 sequences that made the most significant match with DosS CA based on their expect value (<1e^-6^) and aligned their ATP-lids using the COBALT tool. Notably, we observed that the six amino acid lid-length was the most common among the homologous proteins (n=2523), with much fewer proteins exhibiting five (n=7), seven (n=362), eight (n=24), and nine (n=72) amino acids in their lids (**Fig. 4a**). Moreover, only twelve of the top 3000 DosS CA-type proteins displayed a lid with 10-27 amino acids. Next, we evaluated the sequence homology of the 3000 homologous proteins with varied amino acid lid sizes at the N-box, G-box, and ATP-lid residues. A LOGO plot analysis revealed strong conservation of residues E499, N503, R506, and H507 in the N-box (**Fig. 4b**), and D529, G531, and G533 residues in the G1-box (**Fig. 4c**). Furthermore, we found significant conservation of S541 in G2-box and T568 in G3-box of these 3000 homologous proteins (**Fig. S12**). Notably, all the residues that showed H-bonding and ionic interactions in MD simulations of ATP-bound AlphaFold DosS CA model were conserved in a large majority of the 3000 homologous proteins. Additionally, these proteins showed little sequence homology in their ATP-lid regions beyond a conserved proline which could be important for stabilizing/kinking the lid structure (**Fig. S13**). Overall, our bioinformatics studies reveal the ubiquitous presence of short ATP-lids containing less than 10 residues in 2988 bacterial kinases and ATPases and suggests that such features do not impede ATP association in these proteins.

**Figure 4.**
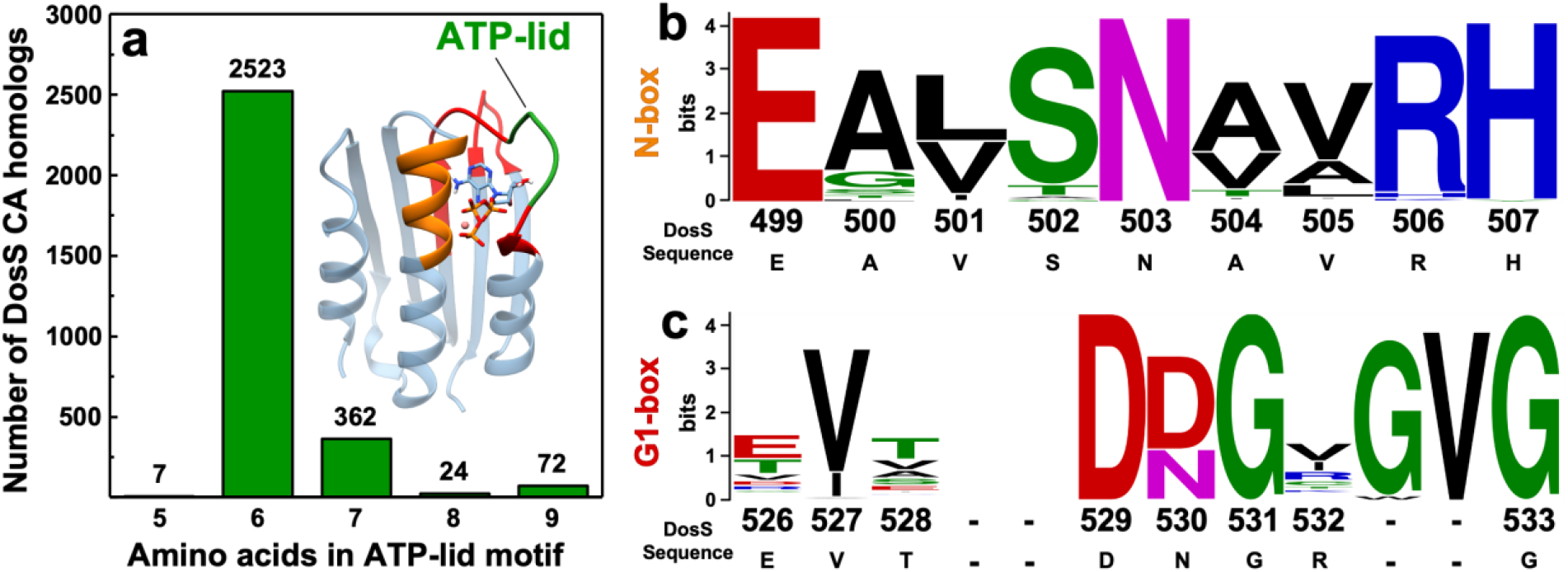
**a)** Bar graph demonstrating the variation in the number of ATP-lid residues in 2988 proteins that are homologous to the DosS CA domain. Logo plots representing the frequency and conservation of amino acids at each position of the (**b**) N box, (**c**) G1 box of 3000 proteins homologous to the DosS CA domain. The height of letter stacks indicate conservation of amino acids at each position, while the height of letters within a stack represents the frequency of an amino acid at each position. Blank spaces in the sequence alignment indicate low conservation of amino acids at these positions. Positions with dashes represent gaps in the DosS CA primary sequence.

## CONCLUSION

Our computational modeling and biophysical studies demonstrate that the ATP pocket in DosS CA is capable of binding ATP with an affinity comparable to other prototypical HKs with F-boxes and longer ATP-lids. MD simulations of DosS CA in the presence and absence of zinc illustrate that the ATP-lid has substantial flexibility in the absence of zinc and that the AlphaFold model is likely providing a physiologically relevant representation of DosS CA as compared to the crystallized zinc-bound structure. Our ITC studies and bioinformatics investigations of homologous proteins further corroborate the MD-based ATP-binding pose obtained via the AlphaFold model. Our findings underscore potential artifacts, such as the random coil transition of an important N-box alpha helix turn in the zinc-bound structure of DosS CA crystallized in the presence of excessive zinc. This highlights the necessity of being cognizant of possible artifacts that can arise in the crystallization of proteins, particularly when parasitic elements employed for crystallization appear at active site positions. Consequently, it conveys the importance of executing comprehensive biophysical, computational, and functional studies to validate the physiological relevance of the static, average structure of proteins obtained from diffraction experiments.

Ultimately, our findings show that DosS CA-like domains are ubiquitous in bacterial species and that ATP binding is possible in CA domains of kinases in the absence of an F-box and a long ATP-lid. DosS CA with an AMP-PNP *K*_d_ of 53 μM, is almost always bound to ATP in bacterial environments where the free ATP concentration is estimated to be 1-5 mM. From a signaling perspective, our studies indicate that ATP binding to DosS CA is unlikely to be a regulatory factor in the auto-phosphorylation reaction and subsequent signaling cascade.

## Author Contributions

GWL performed protein expression, purification, crystallization, CD, ITC studies. GWL and PW performed docking and MD studies. GWL and ES performed bioinformatics studies. KS and HA screened for crystallization conditions and solved the crystal structure. GWL, ARD, and ABD wrote the manuscript with contributions of all authors. GWL, ARD and ABD designed the study. ARD and ABD supervised the study. All authors have given approval to the final version of the manuscript.

## ACKNOWLEDGMENT

This work was supported by the Regents of the University of Minnesota and NIH NIGMS grant #R35GM138277 and R35GM118047. ES was supported by NIH Chemical Biology Training Grant #T32GM132029. X-ray diffraction data were collected at the Northeastern Collaborative Access Team beamlines, which are funded by the NIH (P30 GM124165). The Pilatus 6M detector on 24-ID-C beamline is funded by a NIH-ORIP HEI grant (S10 RR029205).

## ABBREVIATIONS

AMP-PNP: adenylyl imidodiphosphate
BLAST: basic local alignment search tool
CA: catalytic ATP-binding domain
CD: circular dichroism
COBALT: constraint-based alignment tool
DHp: dimerization and histidine phosphotransfer domain
HK: histidine kinase
ITC: isothermal titration calorimetry
*K*_d_: dissociation constant
*Mtb*: *Mycobacterium tuberculosis*
RR: response regulator
TCS: two-component system

## Notes

### Competing Interest Statement

The authors have declared no competing interest.

