## Supplementary Information for "Understanding ATP binding to DosS catalytic domain with a short ATP-lid"

Corresponding Authors

### Table of Contents

#### Supporting Tables:

Table S1. DosS CA crystallization data collection and refinement statistics………...…S3

Table S2. Dissociation constants of nucleotides to ATP-binding proteins……………...S4

#### Supporting Figures:

Figure S1. Generic two component system signal transduction pathway…………...…..S5

Figure S2. COBALT alignment of HK CA domain homology boxes………………..…S6

Figure S3. SDS-PAGE analysis of DosS CA and its variants…………………………….S7

Figure S4. DosS CA with labeled alpha helices and beta strands……………………….S8

Figure S5. ITC titration data of AMP-PNP into DosS CA…………………………..…..S9

Figure S6. ITC titration data of AMP-PNP into DosS CA with 10 nm $Zn(OAc)_2$……..S10

Figure S7. ITC titration data of AMP-PNP into DosS CA with varied $Zn(OAc)_2$………S11

Figure S8. DosS CA crystal, AlphaFold, and homology modeled structures…………..S12

Figure S9. Comparisons of ATP-bound CA domains from various proteins……………S13

Figure S10. Comparing interactions of ATP-bound DosS CA………………………….S14

Figure S11. ITC titration data of AMP-PNP into DosS CA variants……………………S15

Figure S12. Logo plots of the G2 and G3 homology boxes in HK Cas……………..…S16

Figure S13. Logo plots of F boxes in HK CAs………………………………………….S17

**Table S1.** Crystallization data collection and refinement statistics.

| | DosS CA |
|---|---|
| **Data collection** | |
| Space group | C2 |
| Unit cell dimensions | |
| a, b, c (Å) | 54.44, 61.78, 72.23 |
| α, β, γ (º) | 90, 107.54, 90 |
| Resolution (Å) | 34.44 - 1.47 (1.53 - 1.47)* |
| $R_{sym}$ or $R_{merge}$ | 0.034 (0.35) |
| I / σI | 11.63 (1.55) |
| Completeness (%) | 98.38 (93.10) |
| Redundancy | 1.9 (1.8) |
| $CC_{1/2}$ | 0.998 (0.805) |
| **Refinement** | |
| Resolution (Å) | 34.44 - 1.47 (1.53 - 1.47)* |
| No. Reflections | 37912 (3553) |
| $R_{work}$ / $R_{free}$ | 0.153/0.193 |
| No. atoms | 2170 |
| Protein | 1953 |
| Ligand/ion | 37 |
| Water | 180 |
| B-factor | 29.98 |
| Protein | 28.41 |
| Ligand/ion | 46.02 |
| Water | 43.73 |
| R.m.s. deviations | |
| Bond lengths (Å) | 0.004 |
| Bond angles (º) | 0.72 |

*Statistics for the highest-resolution shell are shown in parentheses.

**Table S2.** Dissociation constants of various nucleotide substrates to ATP-binding proteins determined by ITC.

| Protein | Nucleotide | Kd (μM) | Full Length or Domain | Protein Function | DOI |
|---|---|---|---|---|---|
| Protein Kinase A | AMP-PNP | 83 | α subunit of PKA-C | Serine/Threonine Kinase; Regulation of metabolism | 10.1038/s42003-021-01819-6 |
| Protein Kinase A | AMP-PCP | 17.6 | α subunit of PKA-C | Serine/Threonine Kinase; Regulation of metabolism | 10.1126/sciadv.1600663 |
| HK853 | AMP-PNP | 6.8 | Full Length | Histidine Kinase; function not known | 10.1038/ncomms4258 |
| HK853 | ADP | 19 | DHp + CA domains | Histidine Kinase; function not known | 10.3390/molecules24050933 |
| Ks-Amt5 | ATP | 11.2 | DHp + CA domains | Histidine Kinase; Sensing of external ammonium concentrations | 10.1038/s41467-017-02637-3 |
| Ks-Amt5 | ATP-$\gamma$-S | 36.7 | DHp + CA domains | Histidine Kinase; Sensing of external ammonium concentrations | 10.1038/s41467-017-02637-3 |
| Spo0B | ATP | 20 | Full Length | Histidine Kinase; Initiation of sporulation | 10.1111/j.1742-4658.2007.06240.x |
| NifL | ADP | 23.7 | Full Length | Histidine Kinase; Senses redox and fixed nitrogen status | 10.1074/jbc.M610827200 |

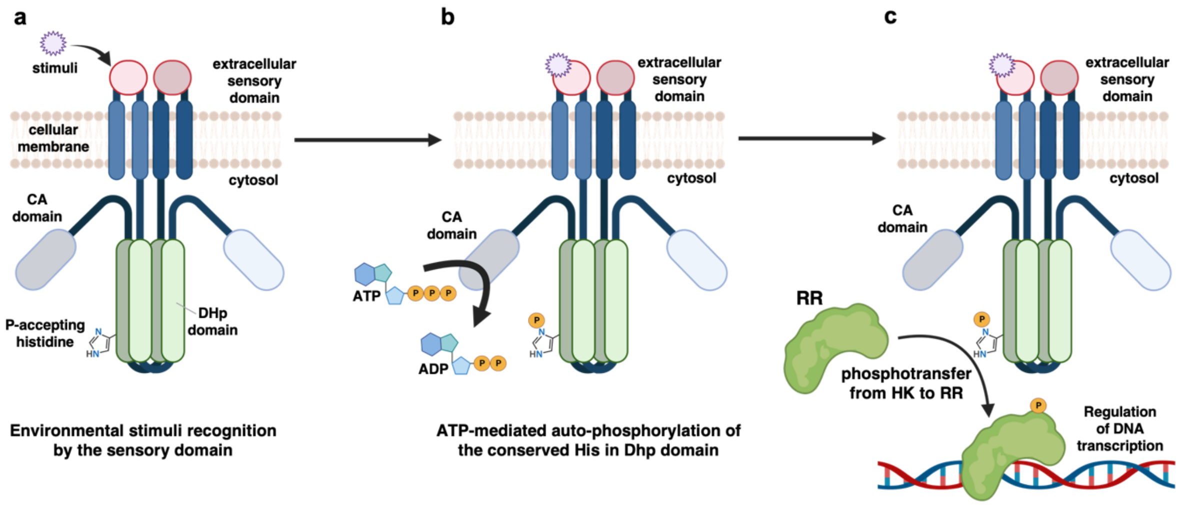


**Figure S1.** A simplistic model of HK and RR-mediated bacterial TCS pathway (**a**) Environmental stimulus is recognized by HK's extracellular sensory domain. (**b**) Stimulus recognition by the sensory domain initiates ATP-mediated auto-phosphorylation of a conserved histidine in DHp domain. (**c**) The signal is further transmitted via phosphotransfer from HK to RR. The phosphorylated RR binds to the DNA and alters gene transcription to allow for cellular adaptation to the environmental stimuli. Note: This model demonstrates a *cis* auto-phosphorylation mechanism in panel b) i.e. CA domain autophosphorylates DHp on the same HK monomer. The mechanism of DosS autophosphorylation is not known.

```
Protein   Species                N box                 G1 box   F box / ATP-lid   G2 box              G3 box

DosS  - M. tuberculosis   - EAVSNAVRHAKASTLTVRVK-VDDDLCIEVTDNGRGLPDEFT------------------GSGLTNLRQRAEQAGGEFTLASVPGASGTVLRWSAPLSQ-----
DosT  - M. tuberculosis   - EAVSNAVRHANATSLAINVS-VEDDVRVEVVDDGVGISGDIT------------------ESGLRNLRQRADDAGGEFTVENMP-TGGTLLRWSAPLR------
WalK  - B. subtilis       - NIISNALKYSEGGHVTFSIDVNEEEEYISVKDEGIGIPKKDV---VFDRFYRVDKARTGTGLGLAIAKEMVQAHGGDIWADSIE-GKGTTITFTLP-YKEEQ--
VraS  - S. aureus         - EAISNTLRHSNGTKVTVELFNKDDYLLLRIQDNGKGFNVDEKLEQ---------------SYGLKNMRERALEIGATFHIVSLP-DSGTRIEVKAPLNKEDS--
DesK  - B. subtilis       - EAVTNVVKHSQAKTCRVDIQQLWKEVVITVSDDGTFKGEENSFSK---------------GHGLLGMRERLEFANGSLHIDTEN---GTKLTMAIPNNSK----
EL346 - E. litoralis      - EVLTNALQHA-SGVVQLRSSVMSGEQRVTVEDDGRGIPEDCDWPN---------------NLGSRIVRQLVQGLGAELNV-TRG-GTGTIVNIDIPLSQQKTLI
EnvZ  - E. coli           - NMVVNAARYG--GWIKVSSGTEPNRAWFQVEDDGPGIAPEQR---LFQPFVRGDSARTGTGLGLAIVQRIVDNHNGMLELGTSE-RGGLSIRAWLPVPVTRA--
PhoQ  - E. coli           - NVLDNACKYC--EFVEISARQTDEHLYIVVEDDGPGIPLSKR---IFDRGQRVDTLRPGQGVGLAVAREITEQYEGKIVAGESM-LGGARMEVIFGRQHSAPKD
CheA  - S. typhimurium    - HLVRNSLDHGVVGNLILSAEHQGGNICIEVTDDGAGLNRERILAKIFAPGFSTAEQVTGRGVGMDVVKRNIQEMGGHVEIQSKQ-GSGTTIRILLPLTLAIL--
FixL  - B. diazoefficiens - NLFRNALEAMR-RELVVTNTPAAD--EVEVSDTGSGFQDDVI-PNLFQTFFTTKDTG--MGVGLSISRSIIEAHGGRMWAESNA-SGGATFRFTLPAADEN---
CpxA  - E. coli           - NIVRNALRYS--TKIEVGFAVDKDG-TITVDDDGPGVSPEDR---IFRPFYRTDEARDGTGLGLAIVETAIQQHRGWVKAEDSP-LGGLRLVIWLPLYKRS---
CckA  - B. abortus        - NLAVNARDAMQ-ITLRTRNIPAADAAVFEVEDTGTGIPADVL-EKIFEPFFTTKEVGKGTGLGLSMVYGIIKQTGGFIYCDSEV-GKGTTFKIFLPRLIEE-KR
```

**Figure S2.** The alignment of N box, G1 box, F box/ATP-lid, G2 box, and G3 box residues in HK CA domains that are structurally homologous to DosS CA. The highly conserved ATP-interacting residues within each box are underlined.

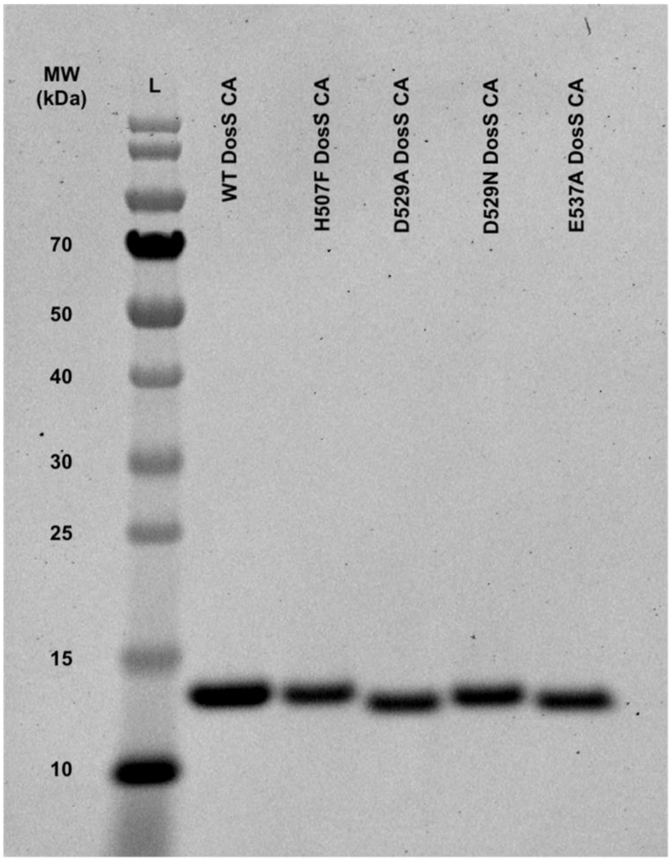


**Figure S3.** SDS-PAGE analysis of WT, H507F, D529A, D529N, and E537A DosS CA variants.

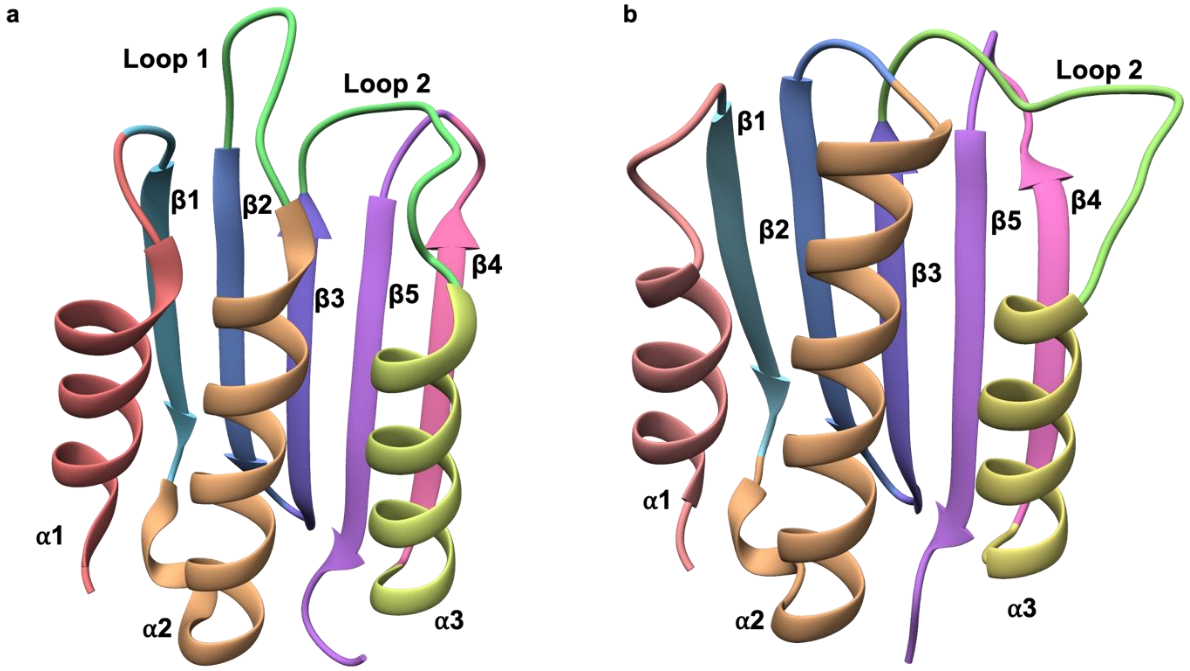


**Figure S4.** (**a**) DosS CA crystal structure with α helices, β strands, and loops colored and labeled (PDB ID: 8SBM). (**b**) DosS CA model with α helices, β strands, and loops colored and labeled.

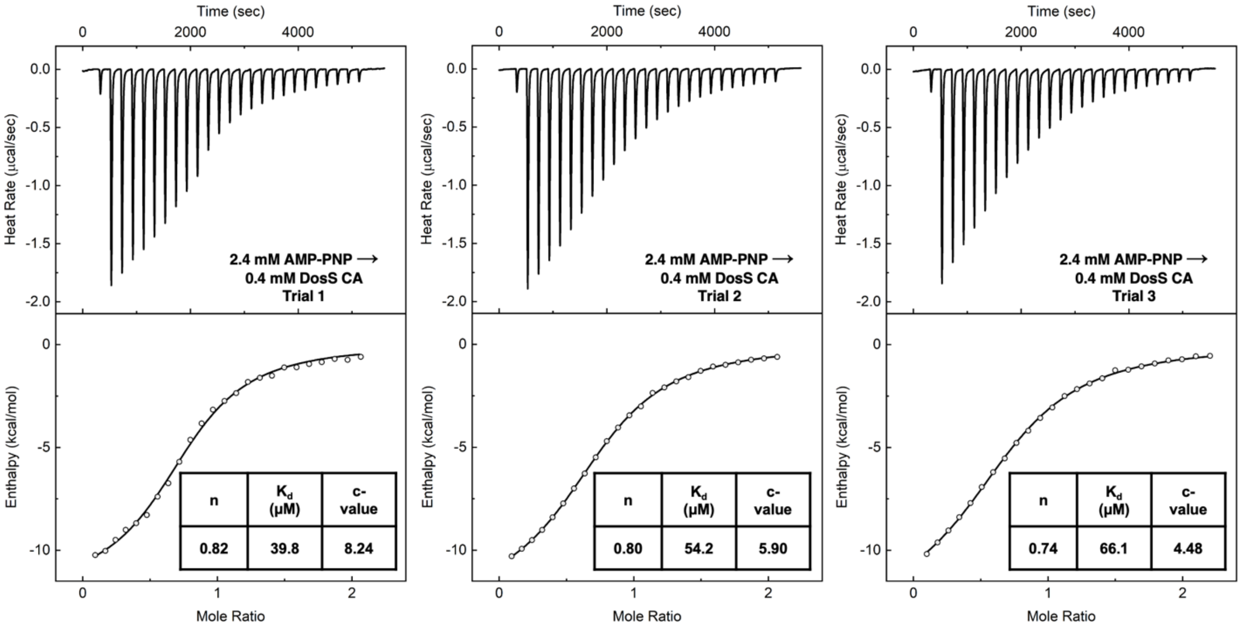


**Figure S5.** Titration data for 2.4 mM AMP-PNP titrations into 0.4 mM DosS CA in triplicate. The top panel of each plot shows the raw heat data collected during the progression of the experiment. The bottom panel represents the integrated heat data plotted against the molar ratio of ligand:receptor.

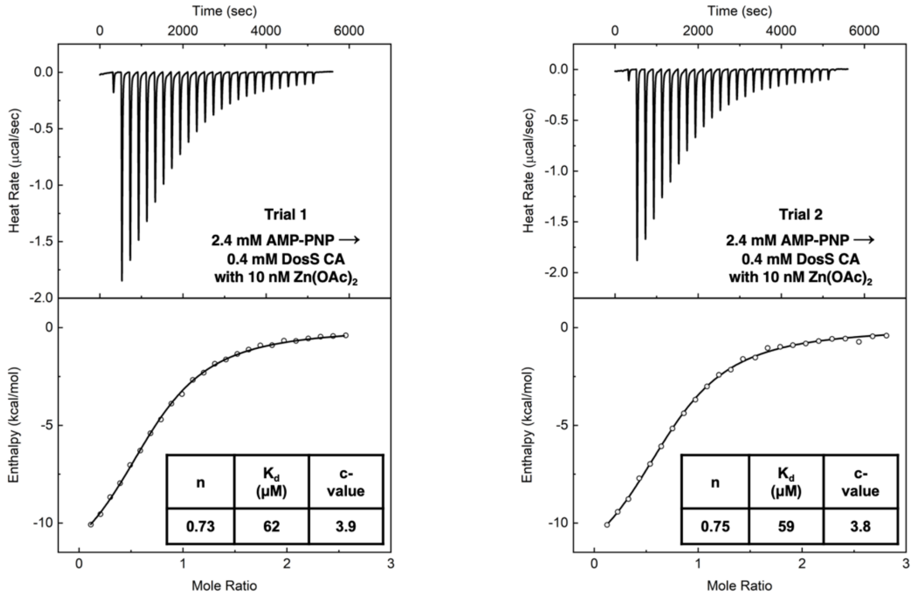


**Figure S6.** Titration data for 2.4 mM AMP-PNP titrated into 0.4 mM DosS CA incubated with 10 nM $Zn(OAc)_2$ performed in duplicate. The top panel of each plot shows the raw heat data collected during the progression of the experiment. The bottom panel represents the integrated heat data plotted against the molar ratio of ligand:receptor.

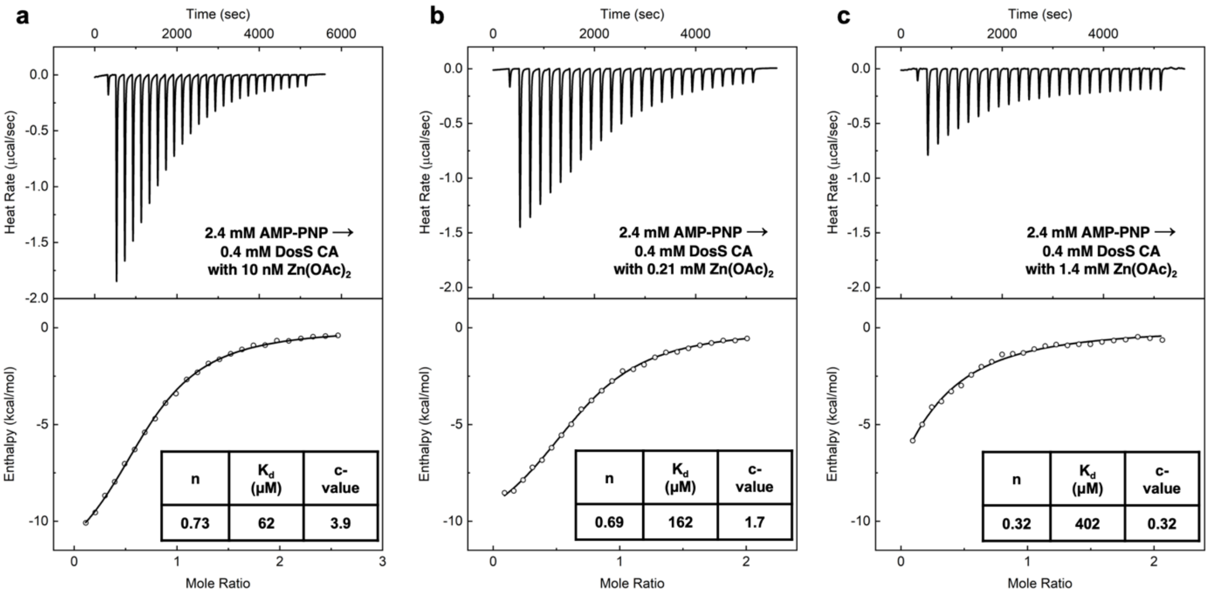


**Figure S7.** Titration data for 2.4 mM AMP-PNP titrated into 0.4 mM DosS CA incubated with (**a**) 10 nM, (**b**) 0.21 mM, and (**c**) 1.4 mM $Zn(OAc)_2$. The top panel of each plot shows the raw heat data collected during the progression of the experiment. The bottom panel represents the integrated heat data plotted against the molar ratio of ligand:receptor.

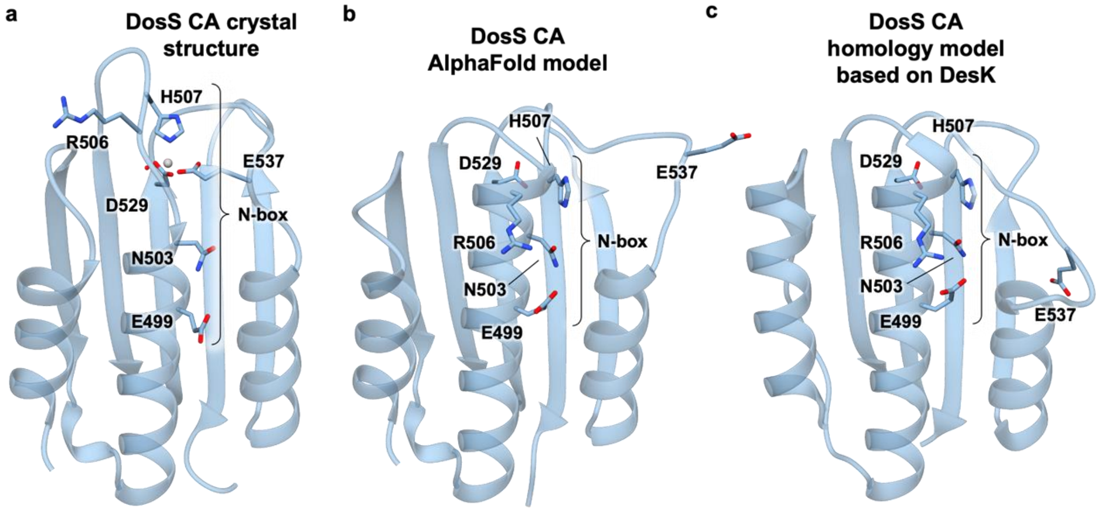


**Figure S8.** (**a**) DosS CA crystal structure (PDB ID: 8SBM). (**b**) AlphaFold predicted DosS CA model. (**c**) DosS CA homology model based on DesK crystal structure (PDB ID: 3EHG).

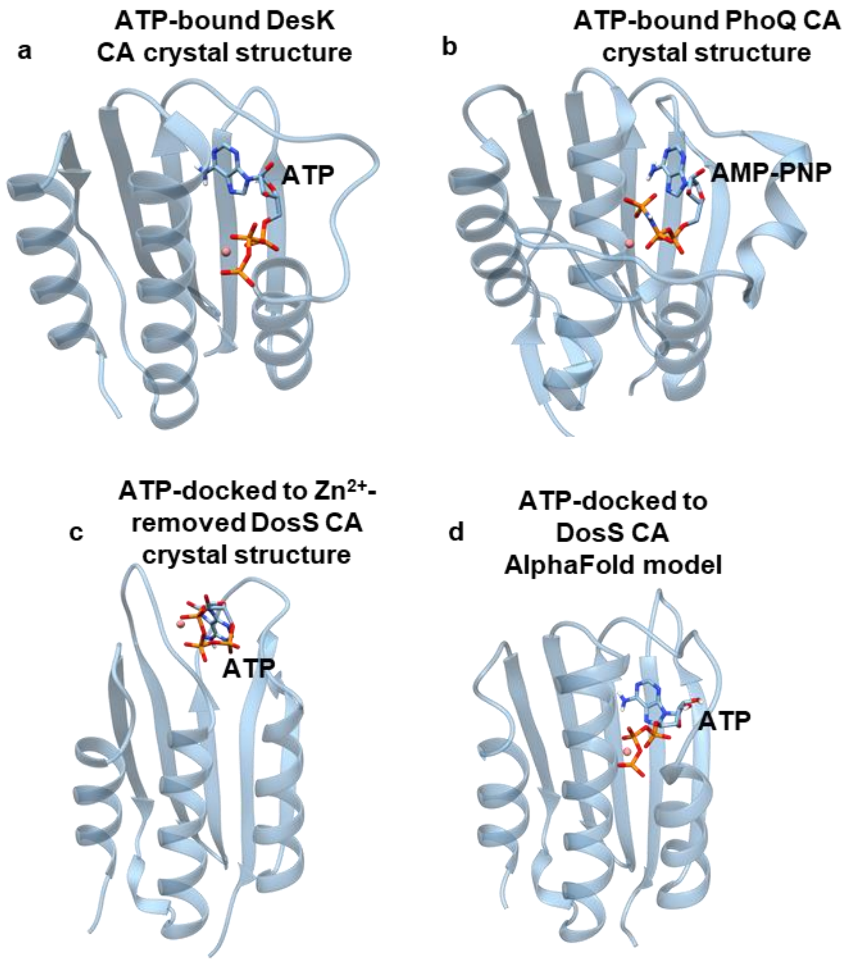


**Figure S9.** Comparisons of ATP-bound CA domains from various proteins. **a)** Crystal structure of DesK CA domain bound to ATP (PDB ID: 3EHG). **b)** Crystal structure of PhoQ CA domain bound to ATP (PDB ID: 1ID0). **c)** ATP docked to an MD simulated DosS CA after removing zinc from the crystal structure. **d)** ATP bound to the AlphaFold predicted structure of the DosS CA domain. The interactions and orientation of ATP in the AlphaFold structure of DosS CA closely resembles that of DesK CA and PhoQ CA structures crystallized with ATP and AMP-PNP, respectively.

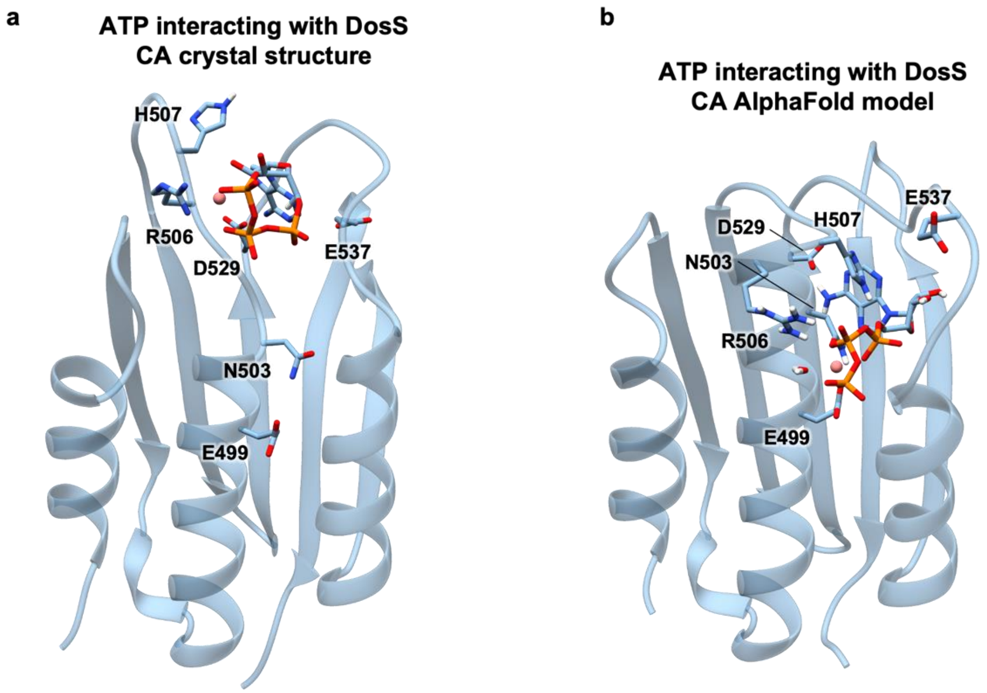


**Figure S10.** (**a**) ATP docked to an MD simulated DosS CA starting from the Zn-bound crystal structure. The distorted alpha helix does not allow for Mg-ATP to interact with all of the N-box residues simultaneously. (**b**) ATP bound to an MD simulated DosS CA starting from the DosS CA AlphaFold model. The intact alpha helix allows for Mg-ATP to interact with all of the N-box residues simultaneously.

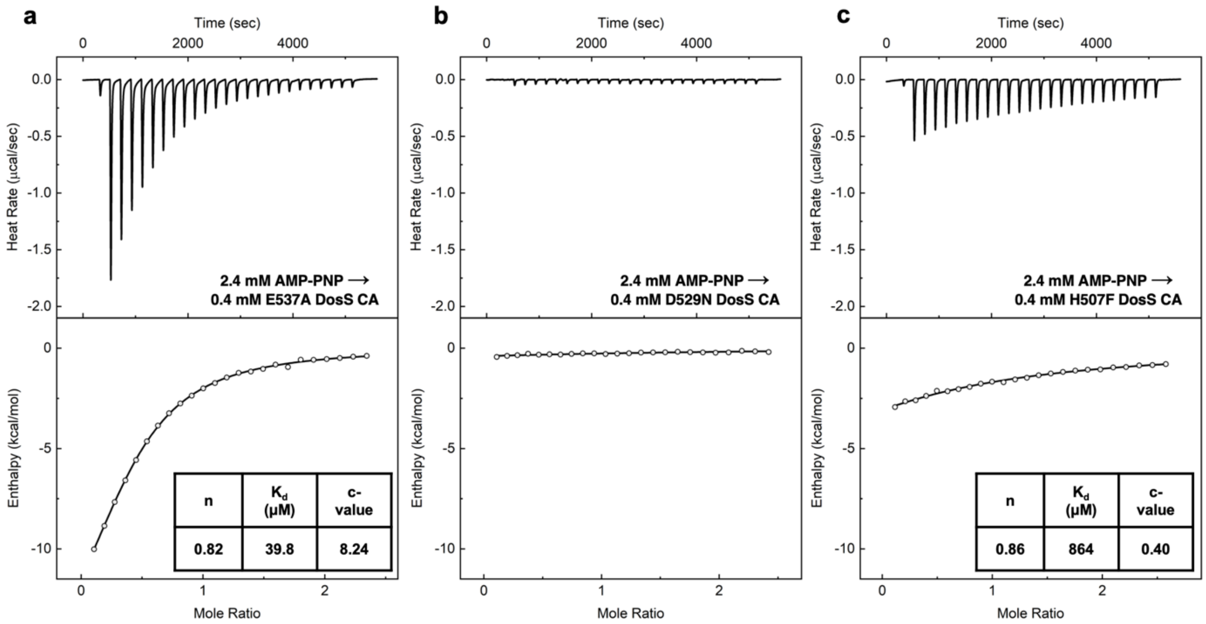


**Figure S11.** Titration data for 2.4 mM AMP-PNP titrations into 0.4 mM (**a**) E537A, (**b**), D529N, and (**c**) H507F DosS CA. The top panel of each plot shows the raw heat data collected during the progression of the experiment. The bottom panel represents the integrated heat data plotted against the molar ratio of ligand:receptor. The c-value for H507F is too low to interpret accurate $K_d$ values.

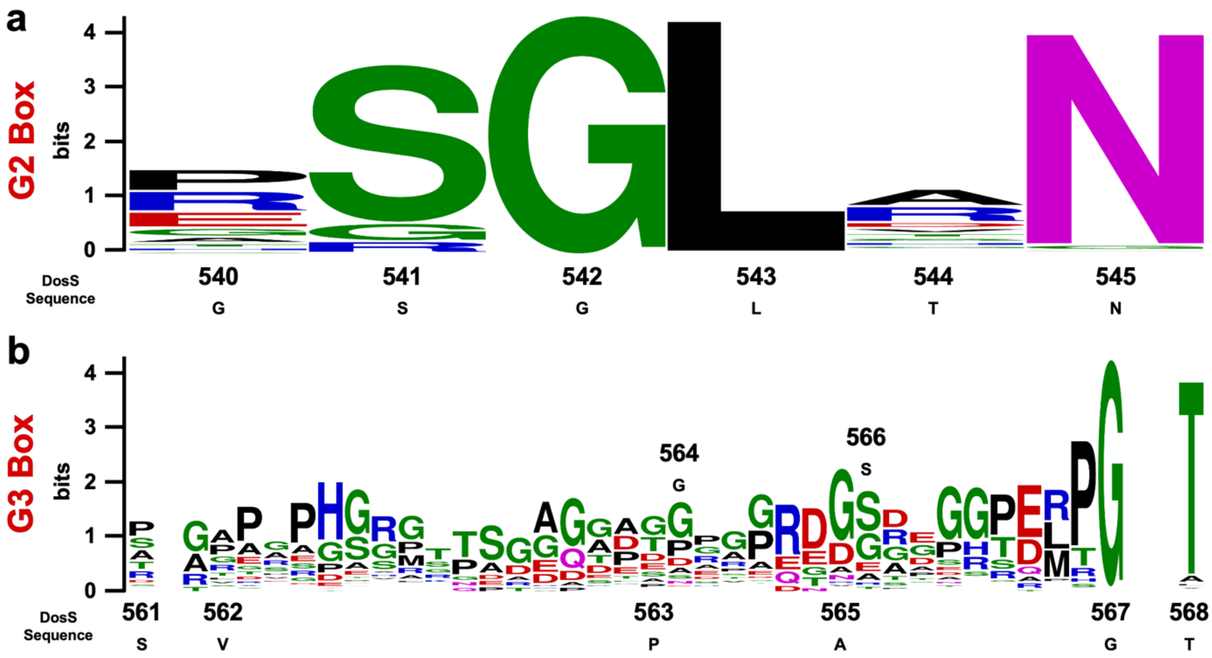


**Figure S12.** Logo plot representing the frequency of amino acids at each position of the (**a**) G2 and (**b**) G3 boxes using the DosS sequence numbering. Positions without labels represent gaps in the DosS CA primary sequence. Blank spaces in the sequence alignment indicate low conservation of amino acids at these positions.

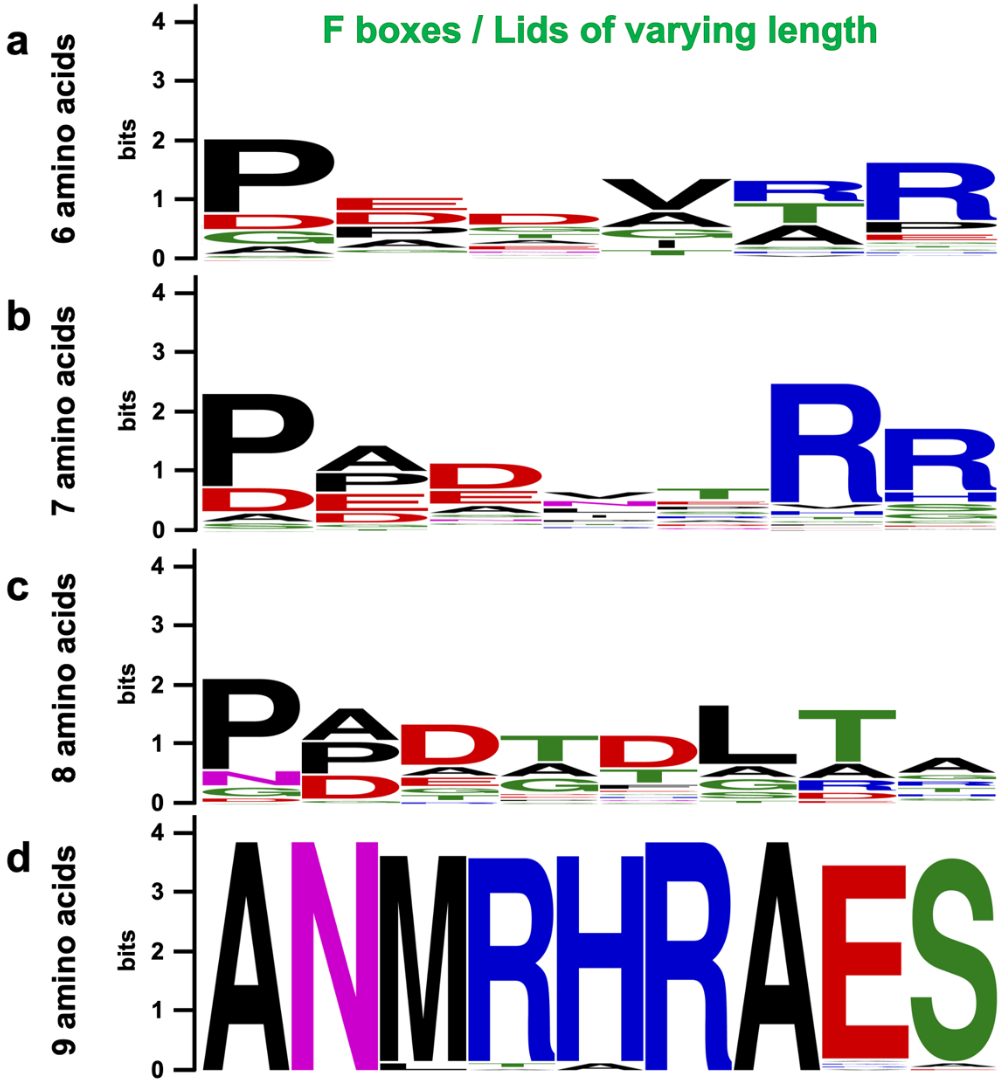


**Figure S13.** Logo plot representing the frequency of amino acids at each position of the F boxes in proteins homologous to the DosS CA. The variation in the number of ATP-lid residues in 2981 proteins that are homologous to the DosS CA domain are represented in logo plots with (**a**) 6 amino acids, (**b**) 7 amino acids, (**c**) 8 amino acids, and (**d**) 9 amino acids.
